# Accurate and Fast Clade Assignment via Deep Learning and Frequency Chaos Game Representation

**DOI:** 10.1101/2022.06.13.495912

**Authors:** Jorge Avila Cartes, Santosh Anand, Simone Ciccolella, Paola Bonizzoni, Gianluca Della Vedova

## Abstract

**Background:** Since the beginning of the COVID-19 pandemic there has been an explosion of sequencing of the SARS-CoV-2 virus, making it the most widely sequenced virus in the history. Several databases and tools have been created to keep track of genome sequences and variants of the virus, most notably the GISAID platform hosts millions of complete genome sequences, and it is continuously expanding every day. A challenging task is the development of fast and accurate tools that are able to distinguish between the different SARS-CoV-2 variants and assign them to a clade.

**Results:** In this paper, we leverage the Frequency Chaos Game Representation (FCGR) and Convolutional Neural Networks (CNNs) to develop an original method that learns how to classify genome sequences that we implement into CouGaR-g, a tool for the clade assignment problem on SARS-CoV-2 sequences. On a testing subset of the GISAID, CouGaR-g achieves an 96.29% overall accuracy, while a similar tool, Covidex, obtained a 77, 12% overall accuracy. As far as we know, our method is the first using Deep Learning and FCGR for intra-species classification. Furthermore, by using some feature importance methods CouGaR-g allows to identify *k*-mers that matches SARS-CoV-2 marker variants.

**Conclusions:** By combining FCGR and CNNs, we develop a method that achieves a better accuracy than Covidex (which is based on Random Forest) for clade assignment of SARS-CoV-2 genome sequences, also thanks to our training on a much larger dataset, with comparable running times. Our method implemented in CouGaR-g is able to detect *k*-mers that capture relevant biological information that distinguishes the clades, known as marker variants.

**Availability:** The trained models can be tested online providing a FASTA file (with one or multiple sequences) at https://huggingface.co/spaces/BIASLab/sars-cov-2-classification-fcgr. CouGaR-g is also available at https://github.com/AlgoLab/CouGaR-g under the GPL.

## 1 Introduction

The global coordination in combating the COVID-19 pandemic has led to the sequencing of one of the largest amount of viral genomic data ever produced. All this data is stored in publicly available archives, such as the European Nucleotide Archive (ENA) and GISAID [15], currently having more than 9.6 million sequenced genomes, classified in *variants, clades*, and *lineages*.

The SARS-CoV-2 virus has evolved since its discovery, and the currently available phylogenies describing its evolutionary history [11] show more than 2000 different genomes, divided into lineages. Since the phylogeny is fairly stable and the main (existing) *lineages*, i.e. the lines of descent, have been identified, a natural and interesting problem is to quickly find, given a sequence, the *clade* to which it belongs, i.e. a group of descendants sharing a common ancestor [1]. Fast and efficient solutions to the clade assignment problem would help in tracking current and evolving strains and it is crucial for the surveillance of the pathogen. This classification problem has been attacked with machine learning approaches [3, 4, 5] using the Spike protein amino acid sequence to drive the classification step.

In this paper we propose a method for classifying SARS-CoV-2 genome sequences based on Chaos Game Representation (CGR) [14]: a deterministic bidimensional representation of a DNA sequence, also called CGR encoding, that can be easily obtained from the genome sequences. The CGR encoding of a sequence has two fundamental properties: it is deterministic, that is there is a unique CGR encoding of each sequence, and reversible, hence the original sequence can be recovered from its representation [6].

A strongly related approach, known as Frequency matrix of Chaos Game Representation (FCGR) [9, 6], starts from the *k*-mers (the substring of length *k*) of the string we want to represent resulting in the the notion of *k*-th order FCGR [36]. The *k-th order FGCR* of a sequence *s* is a *2^k^ × 2^k^* matrix whose elements are the number of occurrences, i.e. the frequencies, of each *k*-mer in s, where each frequency is stored in the specific and distinct position for each *k*-mer. Note that the matrix shape depends on the fact that the sequence *s* is on a 4-symbol alphabet. In essence, the FCGR is an alternative ordering of the histogram for all the *k*-mers (for a fixed integer *k*). Deep Learning and FCGR have been used to evaluate the drug resistance for protein sequences of HIV [23] and for multi-class classification task to identify the source organism for a given protein [10] – in this case the FCGR has been extended to encode sequences in the protein alphabet. The FCGR has also been used for unsupervised clustering of DNA sequences of several species [24] by using dense neural networks, where the input of these networks must be a 1-dimensional vector. In this case, the 2-dimensional FCGR representation of the sequences must be flattened and cannot be fully exploited. For an extensive review on CGR and its applications in bioinformatics, we refer the reader to [21].

Subtyping Sars-Cov-2 sequences has been addressed in the literature with bioinformatics pipelines that require the alignment to a reference genome [11] [35], and also with machine learning approaches aiming to skip the alignment step [26]. Convolutional Neural Networks (CNNs) [19, 20], showed outstanding results in the well-known Imagenet classification problem [18]. To the best of our knowledge, only two works have used CNNs and FCGR for the classification of DNA sequences. In [28] a simplification of the network reported in [20]) was used to classify different taxonomic categories with a dataset of 3,000 sequences (1200–1400 long). A comparison with Support Vector Machines (SVM), showed that CNNs improve over SVM when using a fragment (500bp) of the sequences. In [30], a CNN was proposed for the classification of a dataset of ≈ 660 sequences from eleven phylogenetic families reporting a test accuracy of 87%.

In this paper we leverage the FCGR representation of genomic sequences and CNN power to perform intra-species classification of viral DNA genome sequences, using SARS-CoV2 as our case of study and GISAID clades as our labels. Observe that in this problem the CNN classifies a dataset that is at least two order of magnitude larger than the one considered in the above mentioned papers. Another work that has tackled the clade assignment problem is Covidex [7], a web app tool based on Random Forest and *k*-mer frequencies: to the best of our knowledge this is the most recent work facing our problem. Notice that almost the entire phylogenetics literature deals with inter-species classification, where the distance between possible cluster centroids is larger, hence the classification problem is easier. We propose to use a residual neural network [13] (ResNet50) for the classification of DNA sequences into 11 GISAID clades, using a dataset of two orders of magnitude larger (153K sequences for training) than those analyzed in the above cited works (about 3000 sequences in [28]).

Classification metrics (precision, recall and f1-score) and analysis of the separability of the embeddings generated by the classification layer (Silhouette Coefficient [29], Calinski-Harabasz Score [8], and Generalized Discrimination Value (GDV) [31]) are analyzed for each model. Using the fact that each feature in the FCGR is uniquely related to a *k*-mer, we aim to analyze if the most relevant *k*-mers identified by feature importance methods (Saliency Maps [34] and Shap Values [22]) are related to mutations defining each clade.

We trained four models, one for each value of *k* ∈ {6, 7, 8, 9}. All models performed very similarly, with *k* = 8 being the best one, achieving an overall accuracy of 97.15% in the test set, and the best classification metrics (0.971 for Silhouette Coefficient, 90687.025 for Calinski-Harabasz and −0.734 for GDV). All but two clades (GR and GRY) reported an f1-score above 97% for all the trained models. Since GR is a close ancestor of GRY and these two clades share many mutations, they are confused with each other.

Using the 20 most relevant *k*-mers identified by Saliency Maps, we were able to achieve a similar performance than our CNNs models using SVM for *k* ∈ {6, 7, 8}. Finally, to access the performance of our models w.r.t. other approaches, we compare our results with Covidex [7] the only recent tool that we found in the literature solving the clade assignment problem. Our results show that our models outperform Covidex in all clades and reported metrics (accuracy, precision, recall and f1-score).

## 2 Background

The Chaos Game Representation for encoding DNA/RNA sequences is formally defined as:

#### Definition 1 (Chaos Game Representation (CGR))

*Let s* = *s*_1_… *s_n_* ∈ {*A, C, G, T*}* *be a sequence. Then the CGR encoding of the sequence s is the bidimensional representation of the ordered pair* (*x_n_, y_n_*) *which is defined iteratively as*

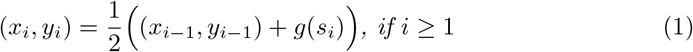

*where* (*x*_0_, *y*_0_) = (0, 0) *and*,

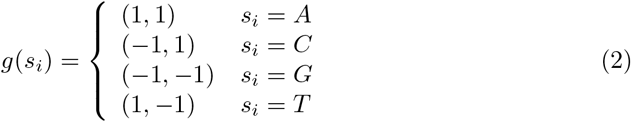

Note that each point (*x_i_, y_i_*) obtained with the above encoding represents the i-long prefix of the sequence *s*. Also, all the CGR encodings are points inside the square with vertices given by the values of the function *g*. In particular, the encoding of all prefixes that shares the last character will be placed in the same quadrant, all prefixes that shares the two last characters, will be placed in the same sub-quadrant, and so on. This property results in a fractal structure of the representation.

Missing bases can be problematic to encode, since the *g*(·) function is not defined in that case, we used the notion of frequence matrix CGR [9, 6], which has the added benefit of allowing us to manage *k*-mers instead of strings of arbitrary length.

### Definition 2 (Frequency matrix of Chaos Game Representation)

*Let s* = *s*_1_… *s_n_* ∈ {*A, C, G, T, N*}* *be a sequence, and let k be an integer. Then the frequency matrix of CGR, in short FCGR, of the sequence s is a* 2*^k^* × 2*^k^ bidimensional matrix F* = (*a_i,j_*), 1 ≤ *i, j* ≤ 2*^k^, i*, 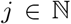. *For each k-mer b* ∈ {*A, C, G, T*}*^k^, we have an element a_i,j_ in the matrix F, that is equal to the number of occurrences of b as a substring of s. Moreover, the position* (*i, j*) *of such element is computed as follows*:

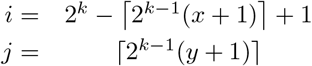

*where* (*x, y*) *is the CGR encoding for the k-mer b*.

Note that the FCGR is defined for a DNA sequence with unknown nucleotide, denoted by *N* — where *k*-mers with an *N* are simply excluded in the counting process—while the *CGR* encoding is well-defined only when all nucleotides are known. To explicitly mention the dimension of the FCGR, we will refer to this as the *k*-th order FCGR.

### 2.1 Classification of viral sequences of DNA

We are given a phylogeny over the possible viral strains, partitioned into classes: each class *c* of such partition 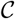 is a *clade* of the tree. More precisely, a clade is a group of related organisms descended from a common ancestor [1], in other words a clade is a subtree of a phylogeny that consist of an ancestral lineage and all its descendants.

Given a genome sequence, that is a string *s* ∈ {*A, C, G, T, N*}*, we determine the original clade in *C* from which the genome sequence is originated; however the genome sequence *s* might not have been previously observed. In any case the sequence will be assigned to a putative clade. To solve this problem, we propose a supervised learning model based on Convolutional Neural Networks (CNN) [20], using FCGR as inputs.

## 3 Data Description

The dataset for this experiment was downloaded from GISAID. By the time of our access to GISAID ^1^ there were around 10 million sequences.

In order to undersample the available data, we first dropped all the rows in the metadata without information in the columns `Virus name, Collection Date, Submission Date, clade, Host` and `Is complete?`, then we built a `fasta_id identifier` from the metadata as a concatenation of the columns `Virus name, Collection Date` and `Submission Date`.

For each clade, we randomly selected 20, 000 sequences considering only those rows where the `Host` column has value ‘‘Human” — clades L, V, and S have less than 20, 000 sequences available, in these cases all sequences have been selected.

As a result of the above procedure, we obtained 191, 456 sequences among the 11 GISAID clades (S, L, G, V, GR, GH, GV, GK, GRY, O, and GRA) over the 12 availables, we excluded the clade GKA from our study since there were only 81 sequences reported in the metadata. The undersampled dataset was randomly split into train, validation and test sets in 80: 10: 10 proportion, preserving the same proportion of clades (labels) in each set. The distribution of the clades over the datasets is given in Table 1

**Table 1:** Distribution of the number of sequences selected for train, validation and test sets by each clade. The final dataset for the 11 clades was split in a 80: 10: 10 proportion for train, validation, and test sets.

| Clade | Train | Val | Test | Total | Available |
| --- | --- | --- | --- | --- | --- |
| S | 14,298 | 1,788 | 1,788 | 17,874 | 17,874 |
| L | 5,154 | 644 | 644 | 6,442 | 6,442 |
| G | 15,999 | 2,000 | 2,000 | 19,999 | 408,552 |
| V | 5,713 | 714 | 714 | 7,141 | 7,141 |
| GR | 16,000 | 2,000 | 2,000 | 20,000 | 625,662 |
| GH | 16,000 | 2,000 | 2,000 | 20,000 | 547,792 |
| GV | 16,000 | 2,000 | 2,000 | 20,000 | 182,248 |
| GK | 16,000 | 2,000 | 2,000 | 20,000 | 4,170,758 |
| GRY | 16,000 | 2,000 | 2,000 | 20,000 | 944,876 |
| O | 16,000 | 2,000 | 2,000 | 20,000 | 55,400 |
| GRA | 16,000 | 2,000 | 2,000 | 20,000 | 2,833,863 |
|  | 153,164 | 19,146 | 19,146 | 191,456 | 9,800,608 |

## 4 Analyses

In this section we present the experimental setup, the dataset used to train and test each model, and clustering and classification metrics. We train one model for each *k* ∈ {6, 7, 8, 9} and we complement the study of the accuracy of each model (compared against Covidex [7]) with an analysis of the most relevant *k*-mers for the classification of each clade using Saliency Maps and Shap.

For this experiment we choose *k* ∈ {6, 7, 8, 9} and sequences from 11 GISAID clades: S, L, G, V, GR, GH, GV, GK, GRY, O and GRA.

### 4.1 Experimental setup

All experiments are conducted using a Intel(R) Core(TM) i5-10400 CPU @ 2.90GHz, x86_64, 32 GB RAM and a graphic card NVIDIA GeForce RTX 3060. The implementation is done in Python 3.9. Tensorflow 2.7.0 [2] was used for training the CNN and scikit-learn 1.0 [25] to compute classification metrics and clustering evaluation (except for Generalized Discrimination Value that was implemented). All code is available online for reproducibility ^2^.

### 4.2 Model training

Each model was set to be trained for 50 epochs with a batch size of 32 (for *k* = 9 we used a batch size of 16 for memory constraints) using Adam optimizer [16] with learning rate 0.001 (the default parameters in keras). The validation loss was monitored after each epoch to save the best trained weights, reducing the learning rate with a patience of 8 epochs and a factor of 0.1, and by an early stopping in case the metrics do not improve after 12 epochs.

Based on the comparison of loss and accuracy of the training and validation sets (see Figures 1 for accuracy and 2 for loss), we observe that training is more stable for *k* = 8 and *k* = 9 in the earlier epochs, i.e. training and validation metrics are similar, while for *k* = 6 and *k* = 7 it takes 33 and 17 epochs to obtain the same behaviour, respectively.

**Figure 1:**
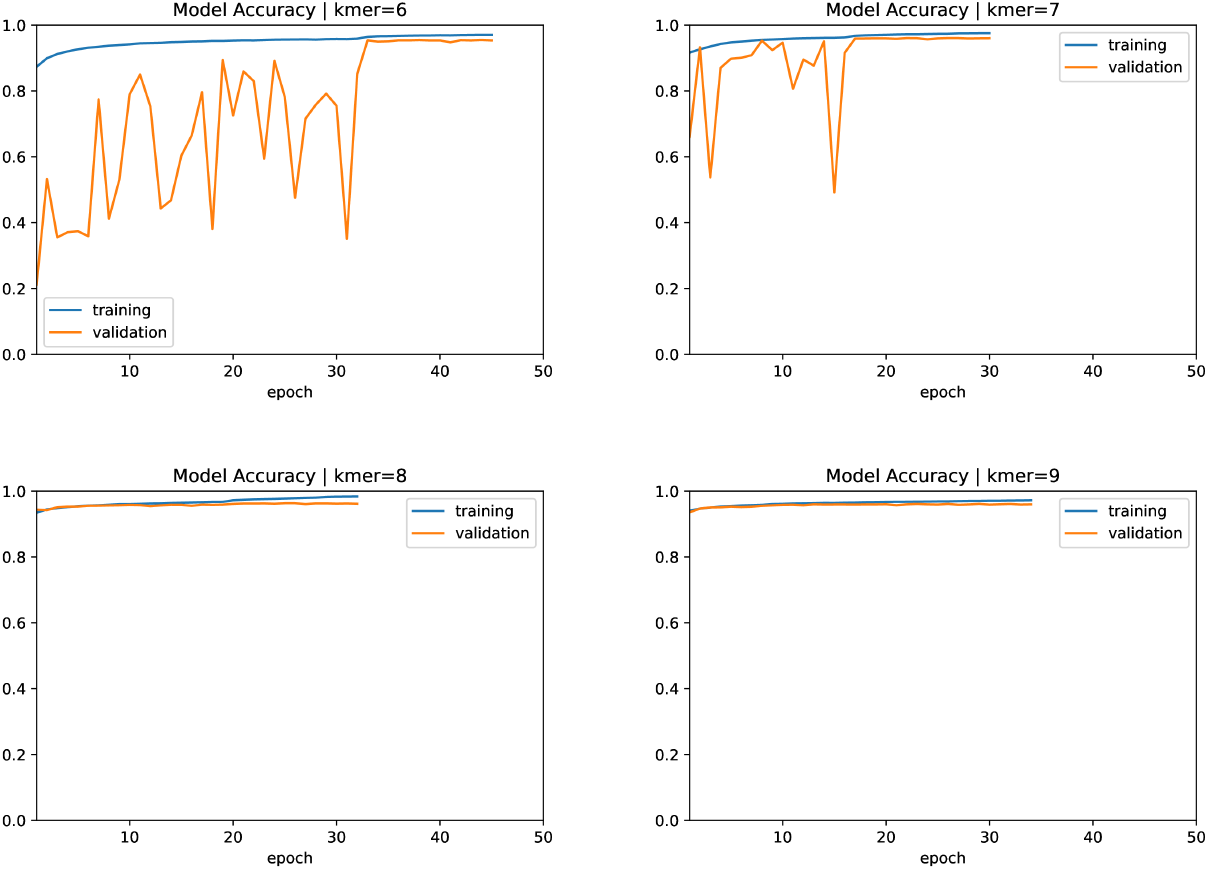
Accuracy in the training and validation sets for our models for *k* ∈ {6, 7, 8, 9}. The models achieve their best performance (based on the validation loss) at epochs 34 (95.4%), 19 (95.9%), 21 (96.1%) and 27 (96.1%), for *k* ∈ {6, 7, 8, 9}, respectively. All models were trained for 50 epochs using an early stopping of 12 epochs based on the validation loss (hence, not all of them ran for 50 epochs). For *k* ∈ {8, 9} the training and validation accuracy are very close, meaning that the model generalizes well in all epochs, while for *k* ∈ {6, 7} it takes several epochs (32 and 15, respectively) to stabilizes.

The architecture used in this experiment is the same for all *k* (ResNet50 [13]), we only changed the input size. Originally, this architecture was designed for inputs of size (224 × 224 × 3), which led us to the assumption that this architecture could be more suitable for *k* = 8. Notice that our sequences are ≈ 29, 000*bp* long, which means that our input FCGR for *k* = 8 is very sparse, since from an n-long sequence we can count *n* – *k* + 1 *k*-mers, it means that (in the case where all *k*-mers are different) we have at most 29, 000 *k*-mers, at least 55% of the elements of the FCGR are 0 for *k* = 8, and a 88% for *k* = 9. In Table 2 a comparison of the number of features for each *k* and the training time per epoch in our experiments is detailed.

**Table 2:** For each *k*, the dimension of the FCGR, its number of features (4*^k^*) and the amount of memory required to store the selected dataset of 191, 456 sequences as FCGR are reported in the table. The number of features and the space increase exponentially w.r.t k.

| $k$-mer | Dimensions | Features | Size (GiB) |
| --- | --- | --- | --- |
| 6 | (64,64) | 4,096 | 6.6 |
| 7 | (128,128) | 16,384 | 24.1 |
| 8 | (256,256) | 65,536 | 94.2 |
| 9 | (512,512) | 262,144 | 374.7 |

### 4.3 Classification results

After each model is trained the precision and recall for the test set are computed for each clade using the best trained weights (lowest loss in the validation set), achieved at epochs 34, 19, 21 and 27 for *k* = 6, *k* = 7, *k* = 8 and *k* = 9, respectively. In our case, we assign each sequence to the clade with highest score. Precision, recall and f1-score are shown in Table 4.

**Table 3:** Accuracy in the test set for each of our models. The model for *k* = 8 exhibits the best accuracy (96.29%), but their accuracy are all in a 1% range.

| $k$-mer | 6 | 7 | 8 | 9 |
| --- | --- | --- | --- | --- |
| Accuracy[%] | 95.45 | 96.08 | 96.29 | 96.17 |

**Table 4:**
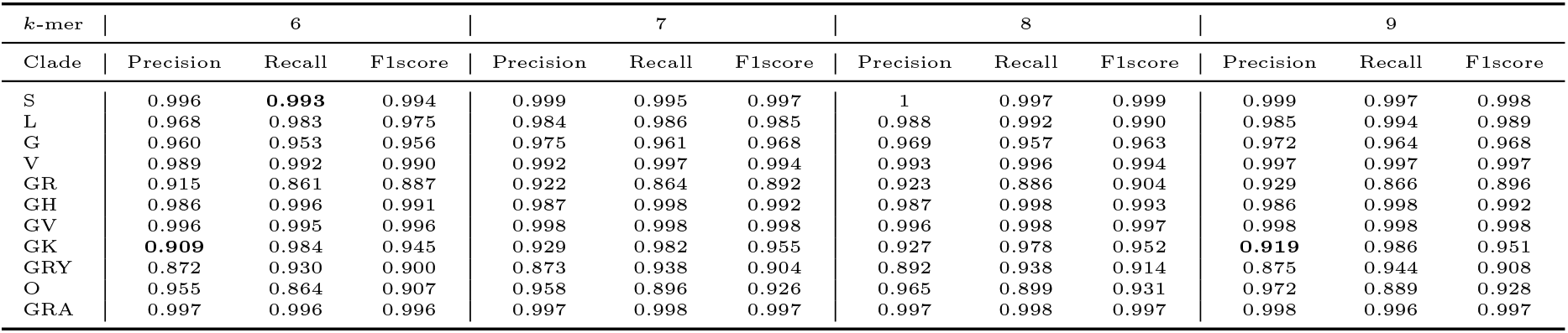
Precision, recall, and f1-score on the test set (19,146 sequences). Each of our models is represented by length of the the *k*-mers used to generate the FCGR. 6 clades out of 11 (S,L,V,GH,GV,GRA) exhibit f1-score above 97.5% in all models. For each clade, precision does not differ more than 2% among the different models (clade GRY between 6-mer and 8-mer). For recall, the largest difference is 3.5% (clade O between 6-mer and 8-mer). Metrics in **bold** are those where **Covidex achieves better performance** than our models in the same dataset.

| *k*-mer | 6 |  |  | 7 |  |  | 8 |  |  | 9 |  |  |
| --- | --- | --- | --- | --- | --- | --- | --- | --- | --- | --- | --- | --- |
| Clade | Precision | Recall | F1score | Precision | Recall | F1score | Precision | Recall | F1score | Precision | Recall | F1score |
| S | 0.996 | **0.993** | 0.994 | 0.999 | 0.995 | 0.997 | 1 | 0.997 | 0.999 | 0.999 | 0.997 | 0.998 |
| L | 0.968 | 0.983 | 0.975 | 0.984 | 0.986 | 0.985 | 0.988 | 0.992 | 0.990 | 0.985 | 0.994 | 0.989 |
| G | 0.960 | 0.953 | 0.956 | 0.975 | 0.961 | 0.968 | 0.969 | 0.957 | 0.963 | 0.972 | 0.964 | 0.968 |
| V | 0.989 | 0.992 | 0.990 | 0.992 | 0.997 | 0.994 | 0.993 | 0.996 | 0.994 | 0.997 | 0.997 | 0.997 |
| GR | 0.915 | 0.861 | 0.887 | 0.922 | 0.864 | 0.892 | 0.923 | 0.886 | 0.904 | 0.929 | 0.866 | 0.896 |
| GH | 0.986 | 0.996 | 0.991 | 0.987 | 0.998 | 0.992 | 0.987 | 0.998 | 0.993 | 0.986 | 0.998 | 0.992 |
| GV | 0.996 | 0.995 | 0.996 | 0.998 | 0.998 | 0.998 | 0.996 | 0.998 | 0.997 | 0.998 | 0.998 | 0.998 |
| GK | **0.909** | 0.984 | 0.945 | 0.929 | 0.982 | 0.955 | 0.927 | 0.978 | 0.952 | **0.919** | 0.986 | 0.951 |
| GRY | 0.872 | 0.930 | 0.900 | 0.873 | 0.938 | 0.904 | 0.892 | 0.938 | 0.914 | 0.875 | 0.944 | 0.908 |
| O | 0.955 | 0.864 | 0.907 | 0.958 | 0.896 | 0.926 | 0.965 | 0.899 | 0.931 | 0.972 | 0.889 | 0.928 |
| GRA | 0.997 | 0.996 | 0.996 | 0.997 | 0.998 | 0.997 | 0.997 | 0.998 | 0.997 | 0.998 | 0.996 | 0.997 |

Precision and recall are very similar among all the trained models, with small improvements when *k* increases, 6 out of 11 clades have f1-score greater than 99% in our best model (*k* = 8). Most notable differences in the performance can be seen in clades GR and GRY, which present the lowest reported recall and precision in each model, respectively. Moreover, from the confusion matrices (see Fig. 3) we can see that misclassified sequences that belong to clades GR and GRY, are confused between them, this can be explained since clade GRY is originated from clade GR. For the other clades, most of the misclassified sequences are predicted as (or belong to) clade G, that is the former one. Clade O exhibits the second lowest recall, where the misclassified sequences are assigned predominantly to clades G, GH, GK, and GRY.

**Figure 2:**
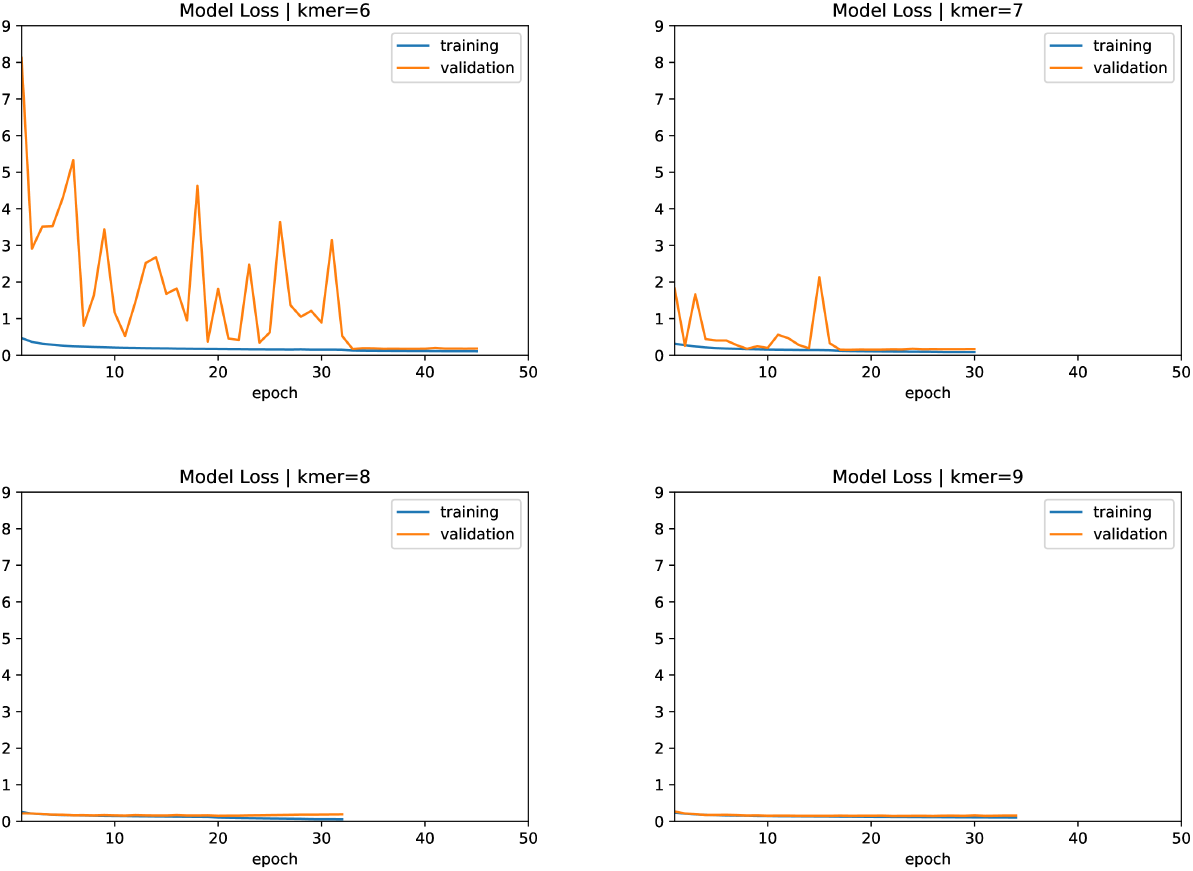
Loss (categorical cross-entropy) in the training and validation sets for the models for *k* ∈ {6, 7, 8, 9}. The best models are chosen as the one with the smallest validation loss. This is achieved at epochs 34 (0.192), 19 (0.156), 21 (0.153) and 27 (0.153), for *k* ∈ {6, 7, 8, 9}, respectively. Models were trained for 50 epochs using an early stopping of 12 epochs based on the validation loss (hence, not all of them run for 50 epochs). Based on the differences between the validation and training losses, the models are able to learn and generalize well in all epochs for *k* ∈ {8, 9}, while for *k* ∈ {6, 7} it takes at least 32 and 15 epochs, respectively.

**Figure 3:**
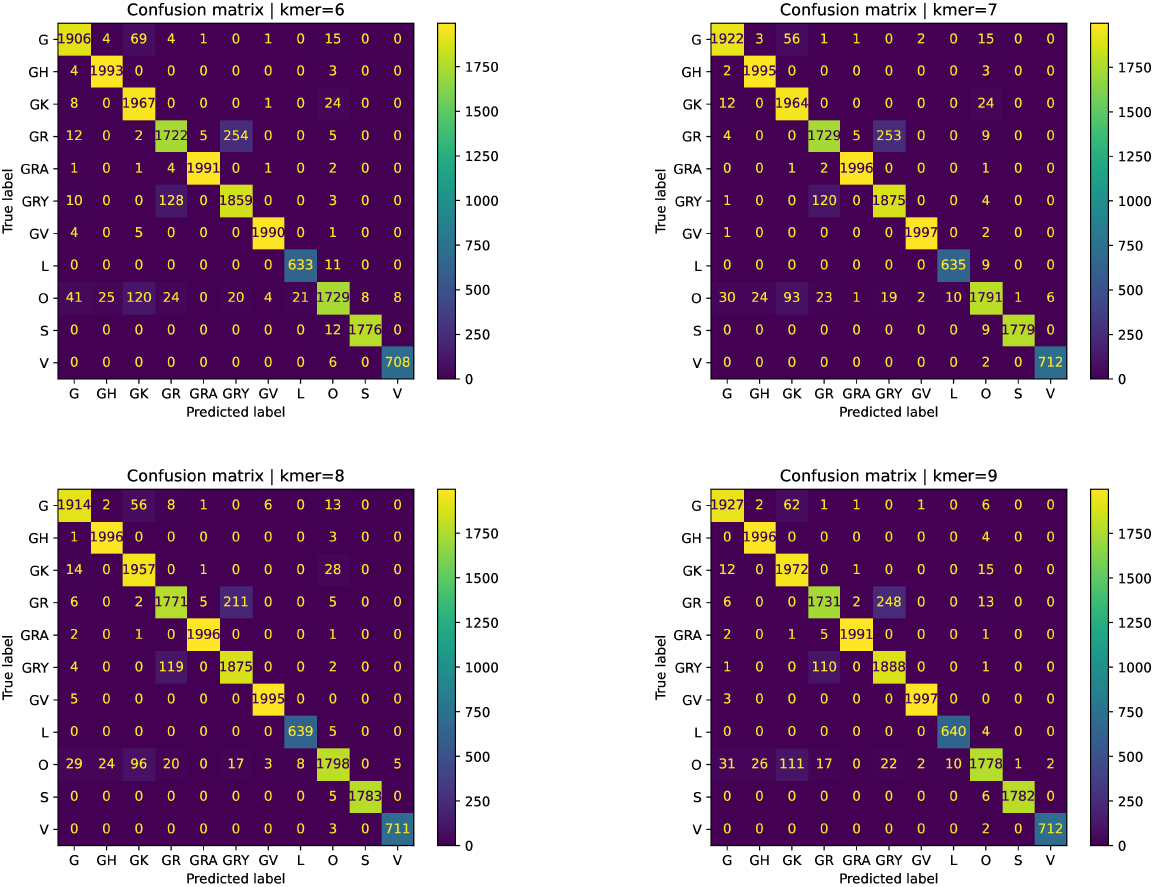
Confusion matrices for the test set for each of our models. All the models are able to correctly classify more than 99% of the sequences for all clades except for G, GR, GRY, and O. Most of the incorrectly classified sequences of GR and GRY are confused between them, which makes sense since they are evolutionary related. For the G clade, the incorrectly classified sequences are shared between clades GH, GK, GR, GV, and O. For the clade O, the incorrectly classified sequences are predominantly assigned to clades G, GH, GK, and GRY.

### 4.4 Comparison with the literature

We compare our results against Covidex [7], a tool that classify Sars-CoV-2 sequences into three nomenclatures: GISAID, Nextrain and Pango lineages. Using a different model for each task, all based on Random Forest and 6-mers as input. The reported accuracy are 97, 77%, 99, 52% and 96, 56% for GISAID, Nextstrain and Pango models, respectively. They also trained the models using 7-mers, but they claim that it only produced slightly better results in terms of accuracy but with more than doubling the computation time [7].

The input for Covidex is a vector with the normalized counting of the frequencies for all 4*^k^ k*-mers. Our input, the FCGR also considers all *k*-mers but in a bidimensional matrix. The main difference between both approaches is the model behind it, while Covidex uses Random Forest to perform the classification, we take advantage of the CNNs and use a 2-dimensional input, the FCGR. Notice that using the FCGR with any other classical Machine Learning method implies to convert the FCGR into a vector, and hence, the loss of the 2-dimensional structure.

Since our model is trained using GISAID clades, we only compare to those results. In Covidex, they used 10 clades: S, L, G, V, GR, GH, GV, GK, GRY and O. In our case, we included GRA since there were enough available sequences by the time of our experiments, but this is not considered in the comparison.

For Covidex, the model for the GISAID nomenclature was trained with 66, 126 sequences and tested on 13, 230. revisionexplain why we did not re-trained Covidex Since Covidex is made available as an user app for any SARS-Cov2 sequence, we used the app over our test dataset to compare the results. We tested Covidex on our test dataset of 17, 146 sequences (excluding the 2000 sequences from GRA clade). Achieving a 77, 12% of accuracy, more than a 18% lower than all our trained models and 20, 65% lower than their reported accuracy. The reported precision, recall and f1-score, as well as the test results over our selected dataset can be seen in Table 5. We found that the reported f1-score of Covidex is quite distant for the one we obtained in our test dataset for clades L (−8.4%), G (−15.8%), GR (−42.4%), GK (−10.9%), GRY (− 19.3%) and O (−28.8%), while for clades S (−0.8%), V (−2.9%), GH (−2.5%) and GV (−0.9%), we can observe a decrement on the reported f1-score ranging from 0.8% to 2.9%. Our models (see Table 4) exhibit better performance than Covidex in all clades and metrics on our test set, with similar results only on clades S and GV. We did not perform an extensive comparison of the running times since both tools classify a genome sequence in less than a second (on *k* = 8, our tool took 0.15 seconds in average).

**Table 5:** Precision, recall, and f1-score for Covidex. The Report part is taken from the Supplementary material of [7]. The Test part has the precision, recall, and f1-score obtained by Covidex on our test set, restricted to the 10 clades (17,146 sequences) analyzed in [7]. We found significant differences between Covidex and our trained models in the Test metrics (see Table 4). In particular, the most notorious differences w.r.t f1-score, ranging from 8.4%–42.4% are found for clades L (−8.4%), G (−15.8%), GR (−42.4%), GK (−10.9%), GRY (−19.3%) and O (−28.8%), while for clades S (−0.8%), V (−2.9%), GH (−2.5%) and GV (−0.9%), we can observe a decrement on the reported f1-score ranging from 0.8%–2.9%. Metrics in **bold** in the Test part are those which **did not decrease** more than 3% w.r.t the reported metrics.

|  | Report |  |  | Test |  |  |
| --- | --- | --- | --- | --- | --- | --- |
| Clade | Prec. | Rec. | F1score | Prec. | Rec. | F1score |
| S | 0.998 | 1 | 0.999 | **0.988** | **0.995** | **0.991** |
| L | 0.997 | 1 | 0.999 | 0.859 | **0.979** | 0.915 |
| G | 0.993 | 0.984 | 0.989 | 0.811 | 0.852 | 0.831 |
| V | 1 | 1 | 1 | 0.958 | **0.985** | **0.971** |
| GR | 0.945 | 0.915 | 0.930 | 0.379 | 0.760 | 0.506 |
| GH | 0.995 | 0.999 | 0.997 | 0.957 | **0.987** | **0.972** |
| GV | 0.996 | 0.999 | 0.997 | **0.980** | **0.995** | **0.988** |
| GK | 0.977 | 0.995 | 0.986 | 0.925 | 0.833 | 0.877 |
| GRY | 0.920 | 0.961 | 0.940 | 0.732 | 0.763 | 0.747 |
| O | 0.994 | 0.959 | 0.976 | 0.722 | 0.658 | 0.688 |

### 4.5 Clustering results

We evaluate the embeddings of the last layer of each trained model using the Silhouette Coefficient, Calinski-Harabasz score and Generalized Discrimination Value (GDV). These results are shown in Table 6. We can observe that the model for *k* = 9 is the best one among all metrics, however, all trained models exhibit a very similar separability based on Silhouette and GDV.

**Table 6:** Clustering metrics for our trained models. Each metric is computed using the output of each model and the predicted clade (that is, the clade that achieves the highest score by our model) in the test set. Each model is represented by the length of the *k*-mers used to generate the FCGR. For the Silhouette score, the closest to 1 the better. For the Calinski-Harabasz score, larger values are better. For the GDV score, the closest to −1, the better. The model with best separability is the one for *k* = 9.

| $k$-mer | Silhouette | Calinski-Harabasz | GDV |
| --- | --- | --- | --- |
| 6 | 0.941 | 150967.217 | -0.713 |
| 7 | 0.944 | 159449.961 | -0.715 |
| 8 | 0.950 | 176844.529 | -0.718 |
| 9 | 0.952 | 183471.005 | -0.720 |

### 4.6 Relevant k–mers for the classification of each clade

Using Saliency Maps and Shap Values, we can evaluate the contribution of each element of a FCGR in the classification, for each model. From each one of these feature attribution methods we can obtain an ordered list of all *k*-mers. For each clade, we use the centroid FCGR of all correctly classified sequences in the test set, then we use each centroid FCGR to identify the most relevant *k*-mers for each clade and then train a SVM using the *N* most relevant *k*-mers (for different values of *N* ∈ {1, 2, 3, 4, 5, 10, 15, 20, 25, 30, 35, 40, 45, 50}) and their respective frequencies as input. The purpose of this experiment is to study if a set of the most relevant *k*-mers (based on feature importance methods) are informative enough to a SVM to perform similarly than the trained CNNs (that uses FCGR as input, and hence all the 4*^k^* possible *k*-mers).

The same training and test sets used for the CNNs were used for the SVM. The results of the accuracy in the test set for the different values of *N* are shown in Fig. 7 6. We can observe that *k*-mers identified by Saliency Maps are more informative than those identified by Shap Values, since for *N* = 20, we obtain similar accuracy in the test set for *k* = 6, 7, 8 compared to CNN (96–97%), while in the case of Shap Values, this accuracy is only achieved by *k* = 7 with *N* = 35. Notice that using *N* = 20, we are considering a small number of all possible *k*-mers (0.49% for *k* = 6, 0.12% for *k* = 7 and 0.03% for *k* = 8).

**Figure 4:**
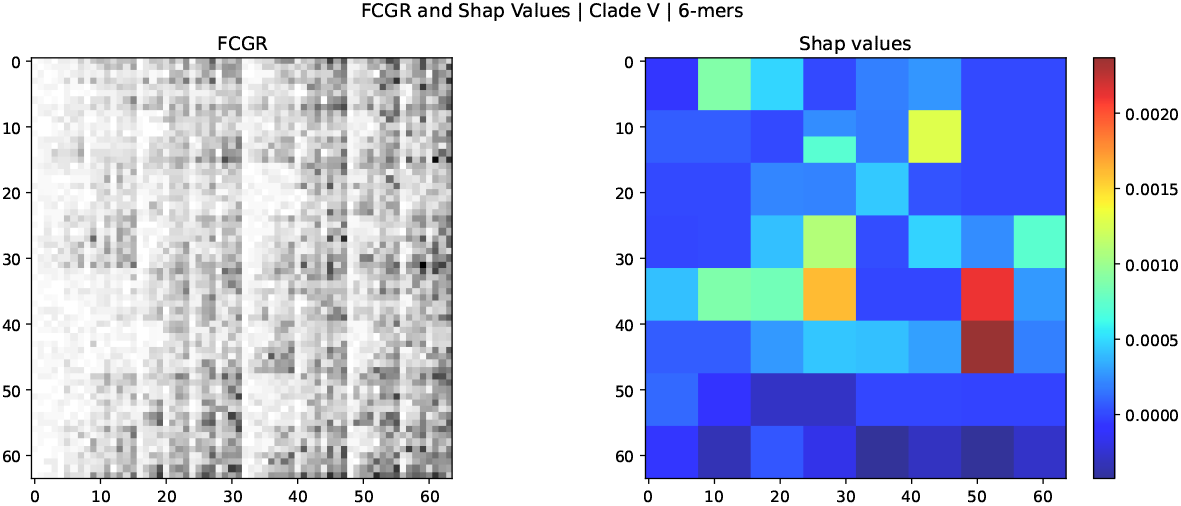
FCGR image (left) and Shap Values (right) of the centroid FCGR for the clade V (*k* = 6). The FCGR image is obtained rescaling the frequencies in the FCGR to a gray-scale range of 8 bits ([0,255]), an inversion of colors is performed to visualize higher values as black squares and lower values as white. Shap Values represent the importance of the features in the FCGR, the higher the value (red) the more important is the feature. Each feature (pixel) in the FCGR corresponds to a *k*-mer.

**Figure 5:**
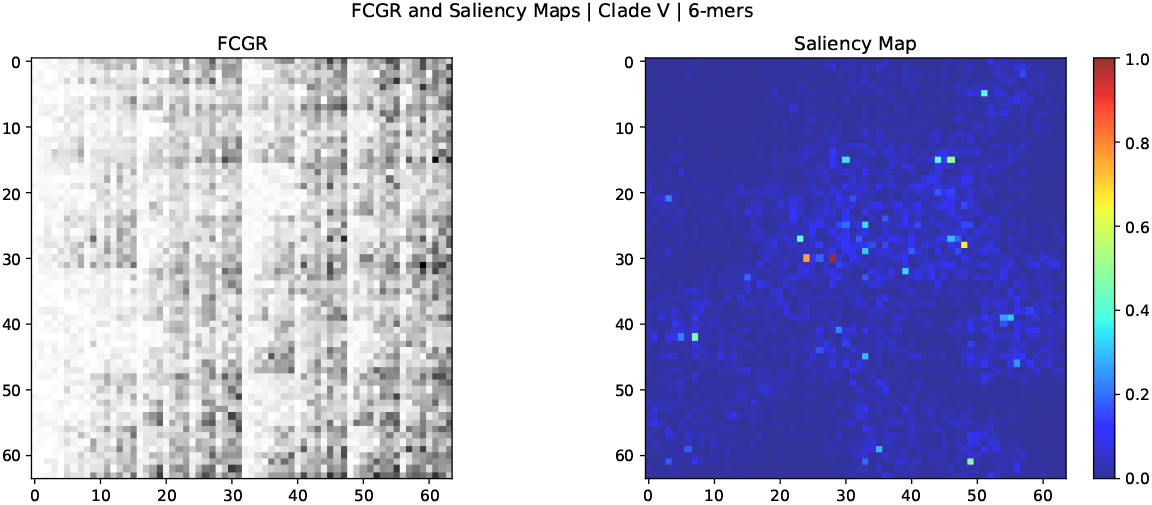
FCGR image (left) and Saliency Map (right) of the centroid FCGR for the clade V (*k* = 6). The FCGR image is obtained rescaling the frequencies in the FCGR to a gray-scale range of 8 bits ([0,255]), an inversion of colors is performed to visualize higher values as black squares and lower values as white. Saliency Map represent the importance of the features in the FCGR, the higher the value (red) the more important is the feature. Each feature (pixel) in the FCGR and Saliency Map corresponds to a *k*-mer.

**Figure 6:**
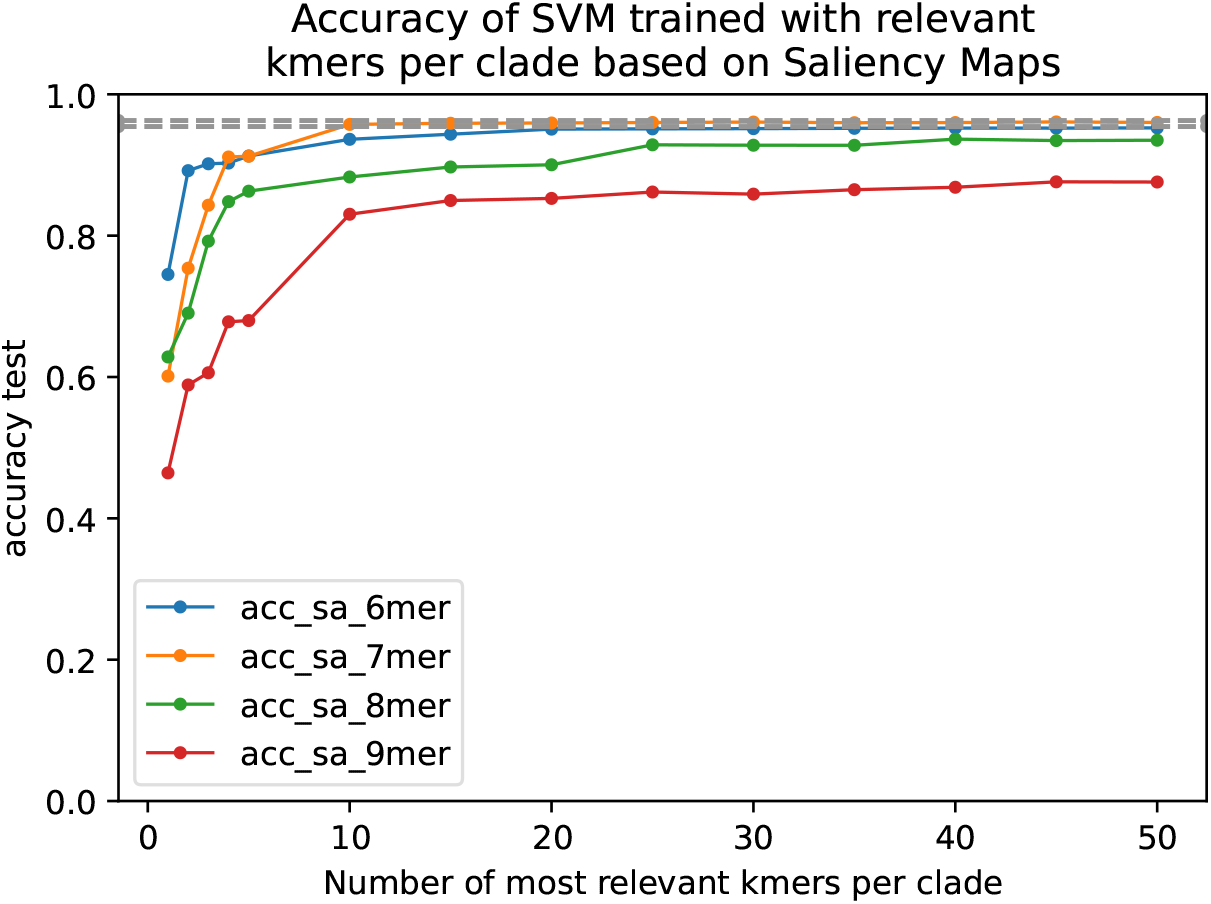
Accuracy of test set for SVM trained models using only the most *N* relevant *k*-mers for each clade (*N* ∈ {1, 2, 3, 4, 5, 10, 15, 20, 25, 30, 35, 40, 45, 50}). The relevant *k*-mers are selected using Saliency Maps on the centroid of the correctly classified FCGR for each clade and model. The same train and test datasets used for the trained CNNs are used for the SVM. The SVM trained with 20 most relevant *k*-mers identified by Saliency Map, for *k* ∈ {6, 7} achieves an accuracy in the test set (≈ 96%) that is in the range of the minimum and maximum accuracies (see 3) obtained by our trained CNNs (the gray dashed band represents the minimum and maximum accuracy for the trained CNNs).

**Figure 7:**
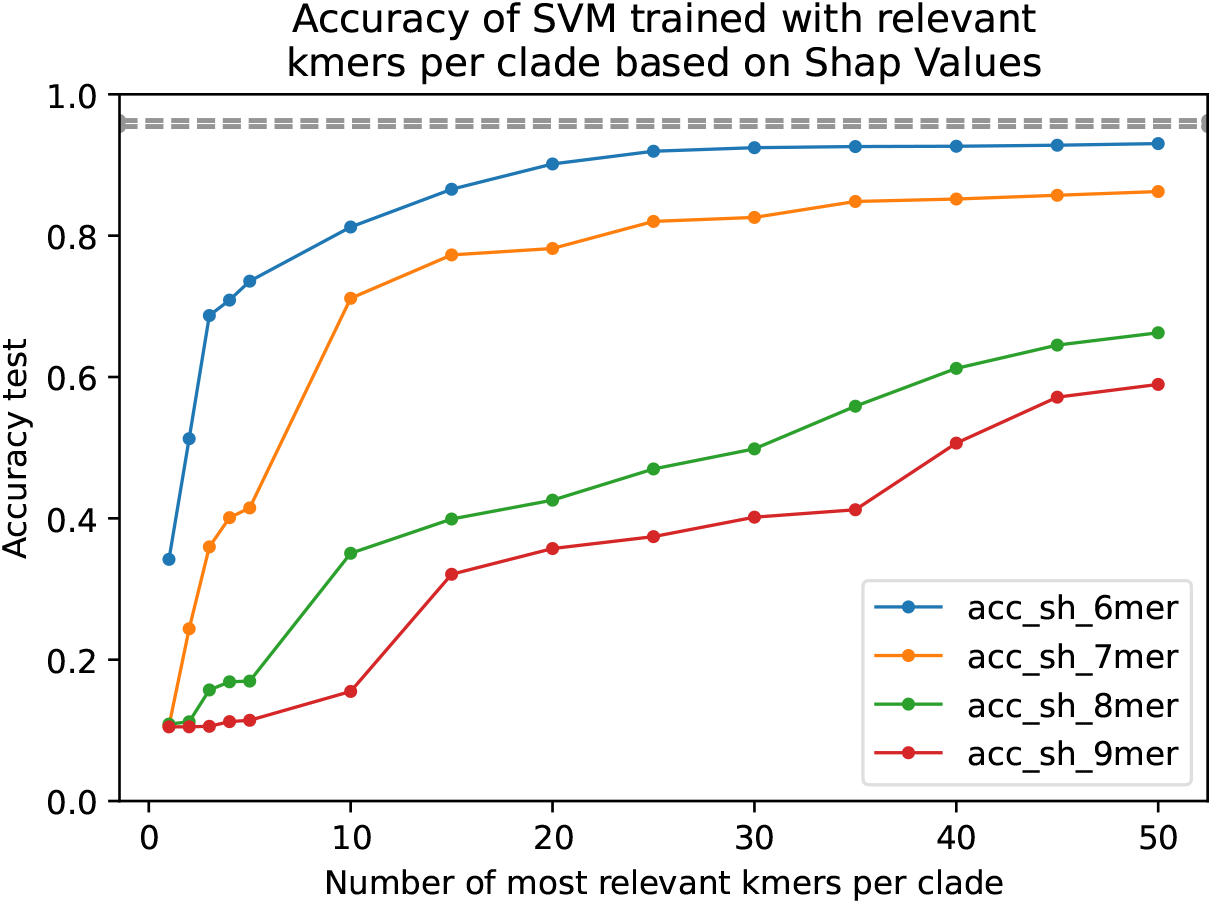
Accuracy of test set for SVM trained model using only the most *N* relevant *k*-mers for each clade (*N* ∈ {1, 2, 3, 4, 5, 10, 15, 20, 25, 30, 35, 40, 45, 50}). The relevant *k*-mers are selected using Shap Values on the centroid of the correctly classified FCGR for each clade and model. The same train and test datasets used for the trained CNNs are used for the SVM. The SVM trained with the 30 most relevant (or more) 6-mers identified by Shap Values, achieves the closest accuracy (92, 44%) to the ones obtained by our trained models (see 3). When *k* increases, the accuracy always decreases (for the same number of relevant *k*-mers), which can be explained since when *k* increases the total number of possible *k*-mers increases exponentially.

### 4.7 Matching relevant k-mers to mutations

Using the reference genome employed by GISAID (EPI_ISL_402124) ^3^ and the list of marker variants ^4^ for each GISAID clade with respect to this reference, we evaluated how many *k*-mers among the 50 chosen ones by Saliency Maps and Shap Values actually matched any of the reported marker variants. A summary is shown in Table 7.

**Table 7:** Summary of matches between the 50 most relevant *k*-mers (from Saliency Maps and Shap Values) and the list of marker variants reported by GISAID for each clade. The *k*-mers obtained by Saliency Maps are able to match several mutations and the matches decrease when *k* increases, but the ones from Shap Values only reported 2 matches, for *k* = 6 and *k* = 7.

| $k$-mer | Saliency Maps | Shap Values |
| --- | --- | --- |
| 6 | 46 | 3 |
| 7 | 51 | 0 |
| 8 | 11 | 0 |
| 9 | 4 | 0 |

The results shown that the most relevant *k*-mers selected using Saliency Maps match several of the reported marker variants (46 matches for *k* = 6, 51 for *k* = 7, 11 for *k* = 8 and 4 for *k* = 9). On the other hand, the ones chosen by Shap Values barely match with the mutation (3 for *k* = 6), suggesting that Saliency Maps could provide a richer explainability of the model from a biological perspective.

## 5 Discussion

In this work we have shown that FCGR can be used to classify DNA sequences. Most notably, we have used FCGR to assign SARS-CoV-2 genome sequences to its GISAID strain by running a CNN on 191, 456 genome sequences (80% training set, 10% validation set, and 10% test set). In particular, the 8-th order FCGR achieved a test accuracy of 96.29%. The majority of missclassified sequences are shared between two strongly related strains, GR and GRY (GR is a close ancestor of GRY).

We have assessed the influence of the length *k* of the substrings (*k*-mers) used to build the FCGR, showing that values between 6 and 9 lead to very similar results, with less than 1% of accuracy on the same test set among them. However, when increasing the value of *k*, the training time for the model and the memory required to save the FCGRs increases exponentially. For *k* = 6 each epoch required 3 minutes and 6.6GB of memory, while for *k* = 9 it required 1:16 hour and 374.7GB. However, FCGRs show a fractal structures; this suggests that we might couple increasing *k* with using only a portion of the FCGR.

We compare our results with Covidex, a Random Forest based tool that classify sequences on GISAID clades based on k-mers frequencies. Under the same test set, our results show that our models outperform Covidex in all clades and reported metrics (accuracy, precision, recall and f1-score). Moreover, we found that the reported precision, recall and f1-score of Covidex are quite different for all clades but S and GV in our test set, exhibiting a decreasing in the f1-score metric up to 42.4%.

We have used Saliency Maps and Shap to identify relevant *k*-mers, looking for matches with the marker variants reported for each strain. Using the *k*-mers obtained by Saliency Map, we found 46, 51, 11 and 4 matches for *k* = 6, 7, 8 and 9, respectively. While, for the *k*-mers identified by Shap, only 3 matches were found for *k* = 6. A possible direction for future works is to explore other existing methods (*e.g*. Lime [27], GradCAM [32], DeepLIFT [33]) that might be suitable in explaining the decisions of the model.

Classifying genome sequences introducing the assembly bias, since any classification depends on the specific assembly pipeline that has been used. To lessen this possible problem, we should study a related problem, where we classify read samples instead of fully assembled genomes. This new problem is more complex, since different regions of the viral genome can have different coverage — hence impacting the frequencies — and reads needs to be cleaned from both errors and contamination artifacts (the latter might be attacked with specialized tools like KMC3 [17].

We did not perform an extensive comparison of the running times since both tools classify a genome sequence in less than a second.

## 6 Methods

We use the *k*-th order FCGR representation for each sequence. In order to obtain this representation, we need to count the *k*-mers in each sequence and to put those frequencies in the FCGR based on the CGR encoding.

Before feeding the FCGR to the model, we rescale its elements to values between 0 and 1 for stability of the learning process. To do so, we divide each FCGR element-wise by the maximum value in the FCGR. It is worth mentioning that other preprocessing steps were taken into consideration but were ultimately excluded because found empirically worse.

### 6.1 Model architecture

We choose a residual neural network, Resnet50 [12] as our CNN, adapted for *k*-th order FCGR, i.e. with input size equal to (2*^k^* × 2*^k^* × 1), and output size equal to the number of clades: 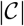, with softmax activation function in the last layer and categorical crossentropy as loss function, since we want to assign only one clade to each DNA sequence.

### 6.2 Model evaluation

To assess the performance of our trained model, we perform a classification evaluation of the predictions and also a clustering evaluation for the embeddings in order to evaluate the class separability.

#### 6.2.1 Classification metrics

We report two classification metrics for each clade in the test set: precision and recall. Given a clade *c*, the correct predictions of the model can be compared to all the sequences with ground truth *c* (recall), and to all the sequences predicted by the model into the clade *c* (precision).

Formally, given a clade *c*, the positive class *P* consists of the set of genome sequences that are assigned to *c*, while all other genome sequences are the negative class *N*. Consequently, the true positive consist of the sequences that originate from the clade *c* and have been assigned to *c*, the false positive consist of the sequences that do not originate from the clade *c* and have been assigned to *c*, the false negative consist of the sequences that originate from the clade *c* and have not been assigned to *c*. The precision and recall are computed as follows:

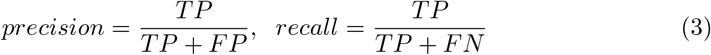

We also report the f1-score, defined as,

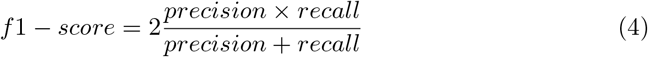

#### 6.2.2 Clustering measures

In order to assess the quality of the class separability given by the CNN, we evaluate the embeddings of the last layer (the one used to perform the classification) in the network with three clustering evaluation measures. These embeddings are the output from the final layer of the network for each FCGR.

##### 1. Silhouette Coefficient

[29] Given an embedding v belonging to a cluster *A*, the silhouette coefficient *s*(*v*) of *v* compares the mean intra-cluster distance in *A* (*a*) with the mean nearest-cluster distance for *v* (*b*), that is, the closest cluster to *v* different from *A*.

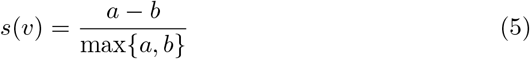

where 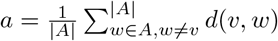 and 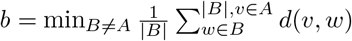.

The value of *s*(*v*) ranges between −1 (wrongly assigned) and 1 (perfect separability). For a cluster *A*, the mean silhouette coefficient of *A* is computed as the average of *s*(*v*) over all embeddings *v* ∈ *A*.

##### 2. Calinski-Harabasz Score

[8] Given a set of embeddings *E* of size *n_E_* that has been clustered into *k* clusters, the Calinski-Harabasz Score *s*, also known as the Variance Ratio Criterion, is defined as the ratio of the between-clusters dispersion and the inter-cluster dispersion for all clusters (the dispersion of a group of *n* points is measured by the sum of the squared distances of the points from their centroid).

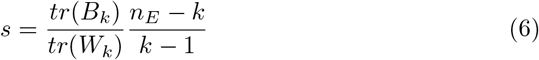

where *tr*(*B_k_*) is the trace of the between-cluster dispersion matrix and *tr*(*W_k_*) is the trace (the sum of all elements in the diagonal of *W_k_*) of the within-cluster dispersion matrix, defined as follow:

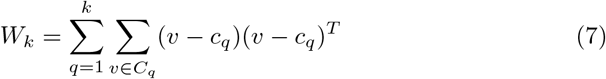

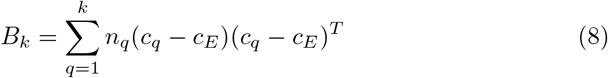

where *C_q_* is the set of embeddings in the cluster *q, c_q_* is the centroid of the cluster *q, c_E_* is the centroid of *E* and *n_q_* = |*C_q_*|.

The higher the score *s* means that the clusters are dense and well separated.

##### 3. Generalized Discrimination Value (GDV)

[31] Given a set of *N* D–dimensional embeddings {*x*_1_,…, *x_N_*}, with *x_n_* = (*x*_*n*,1_,…, *x_n,D_*) and a set of *L* classes {*C*_1_,…, *C_L_*}, where each *x_n_* is assigned to one of the *L* distinct classes. Consider their z–scored points (*s*_1_,…, *s_N_*), with *s_i_* = (*s*_*i*,1_,…, *s_i,D_*), where 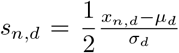. Here 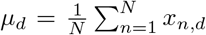 denotes the mean, and 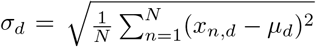 the standard deviation of dimension d. Using the rescaled data points *s_n_* = (*s*_*n*,1_,…, *s_n,D_*), the Generalized Discrimination Value Δ is calculated from the mean intra-class and inter–class distances as follows:

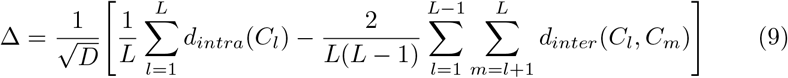

where the mean intra–class for each class *C_l_* is defined as

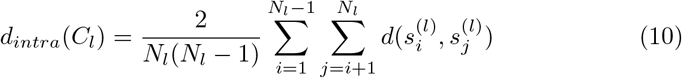

and the mean inter–class for each pair of classes *C_l_* and *C_m_* is defined as follows,

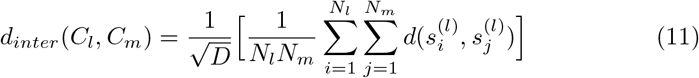

here *N_k_* correspond to the number of points in class *k*, and 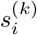 is the ith point of class *k*. The quantity *d*(*a, b*) is the distance between *a* and b, for our case, we considered the Euclidean distance. The value Δ range between −1 (perfect separability) and 0 (wrongly assigned),

### 6.3 Feature importance

After the model is trained, we can perform feature importance methods (also known as pixel attribution in case of images) to analyze the impact of each element of the FCGR in our prediction. We selected Saliency Maps [34] and Shap Values [22]. Saliency Maps calculate the gradient of the loss function for a specific desired class with respect to the input (FCGR) elements, the gradients are rescaled to [0, 1], where elements with values closer to 1 represent the more influential features for the input FCGR over the predicted class. Shap (Shapley Additive Explanations) Values is a game theoretic approach to explain the output of any machine learning model. It aims to explain the influence of each feature compared to the average model output over the dataset the model was trained on, it outputs positive and negative values, where positive values push the prediction higher, and negative values push the prediction lower. Using the most relevant features from both methods over the FCGR, we aim to identify the most relevant *k*-mers for the classification of each clade.

Using these methods we aim to analyze the most relevant *k*-mers for the classification of each clade in the trained models.

## 7 Funding

This project has received funding from the European Union’s Horizon 2020 Innovative Training Networks programme under the Marie Skłodowska-Curie grant agreement No. 956229.

This project has received funding from the European Union’s Horizon 2020 Research and Innovation Staff Exchange programme under the Marie Skłodowska-Curie grant agreement No. 872539.

## 8 Acknowledgements

The authors would like to thank Yuri Pirola, Raffaella Rizzi, Luca Denti, Murray Patterson, and Sarwan Ali for many useful discussions on the topic.

## Footnotes

1 April 04, 2022. https://www.gisaid.org/

2 https://github.com/AlgoLab/fcgr-cnn

3 https://www.gisaid.org/resources/hcov-19-reference-sequence/

4 https://www.gisaid.org/resources/statements-clarifications/clade-and-lineage-nomenclature-aids-in-genomic-epidemiology-of-active-hcov-19-viruses/

## Notes

### Competing Interest Statement

The authors have declared no competing interest.

